# CellAgent: LLM-Driven Multi-Agent Framework for Natural Language-Based Single-Cell Analysis

**DOI:** 10.1101/2024.05.13.593861

**Authors:** Yihang Xiao, Jinyi Liu, Yan Zheng, Shaoqing Jiao, Jianye Hao, Xiaohan Xie, Mingzhi Li, Ruitao Wang, Fei Ni, Yuxiao Li, Zhen Wang, Xuequn Shang, Zhijie Bao, Changxiao Yang, Jiajie Peng

## Abstract

Single-cell RNA sequencing (scRNA-seq) and spatial transcriptomics (ST) data analysis are pivotal for advancing biological research, enabling precise characterization of cellular heterogeneity. However, existing analysis approaches require extensive manual programming and tool manipulation, posing significant challenges for researchers. To address this, we introduce CellAgent, an autonomous, LLM-driven approach that performs end-to-end scRNA-seq and spatial transcriptomics data analysis through natural language interactions. CellAgent employs a multi-agent hierarchical decision-making framework, simulating a “deep-thinking” workflow to ensure that each analytical step remains consistent with the overall task objective. To further enhance its capabilities, we developed sc-Omni, a high-performance, expert-curated toolkit that consolidates essential tools for scRNA-seq and spatial transcriptomics analysis. Additionally, we introduce a self-reflective optimization mechanism, enabling automated, iterative refinement of results through specialized evaluation methods, effectively replacing traditional manual assessments. Benchmarking against human experts demonstrates that CellAgent achieves approximately 60% improvement in efficiency across multiple downstream applications. In terms of accuracy, it maintains performance comparable to existing approaches while preserving natural language interactions. By translating natural language interactions into optimized analytical workflows, CellAgent establishes a scalable paradigm for LLM-driven scientific discovery, bridging the gap between experimental biologists and complex data analytics. This framework minimizes reliance on manual coding and exhaustive deliberation, ushering in the era of the “AI Agent for Science.”

## 1 Introduction

In recent years, the rapid development of technologies such as single-cell RNA sequencing (scRNA-seq) and spatial transcriptomics (ST) has revolutionized molecular biology, enabling scientists to investigate biological systems with unprecedented precision and depth [1, 2]. These biological technologies have produced many large-scale datasets that require advanced computational tools to extract meaningful biological information efficiently. Therefore, numerous computational tools have been developed, including statistical methods [3–6], deep learning methods [7–9], and methods based on large foundation models [10–12], thereby driving innovation and progress in molecular biology. Although these tools have significantly improved the efficiency and accuracy of scRNA-seq/ST data analysis, the manual and flexible application of various tools to analyze complex and massive single-cell sequencing data carries high labor costs. For example, researchers need to carefully choose the appropriate tools, while configuring appropriate hyperparameters that are suited to the unique characteristics of the input data [13]. To effectively conduct such analyses, researchers must possess advanced programming skills along with a strong foundation in biology, which increases the complexity and cost of performing single-cell data analysis tasks [14]. This creates a significant barrier that inhibits the discovery of biological mechanisms. Therefore, there is an urgent need for a more automated, natural language interaction-based, and functionally integrated analysis tool, which may enable users to perform the analysis by “chat”. It can reduce technical barriers, improve data processing efficiency, and further promote knowledge discovery with single-cell RNA-seq and spatial transcriptomics technologies. Recently, with the impressive capabilities demonstrated by large language models (LLMs) [15–19], LLM-driven autonomous AI agents have been developed to automate a range of tasks, such as ChatMOF [20] and ChemCrow [21].

Inspired by these advancements, a new question arises: Can we leverage LLMs to create a biologically proficient LLM-driven agent framework for automating scRNA-seq and ST data analysis tasks?

Although LLMs such as GPT-4 and several LLM-driven agents such as AutoGen [17] can generate codes for scRNA-seq and spatial transcriptomics data analysis, they still fail to address several key challenges. Existing tools are unable to automatically evaluate the output of the analysis, as the evaluation process involves not only assessing whether the code runs correctly but also determining its biological accuracy. Without automatic evaluation, it becomes challenging to select appropriate algorithms and optimize parameters for different datasets in an automated manner. Unfortunately, existing studies indicate that specialized optimization is necessary for different datasets [1, 22]. On the other hand, LLM-driven agents [17–19, 23] typically use LLMs as the core “brain” to decompose complex tasks, call external tools, and coordinate various actions, all of which require comprehensive and accurate knowledge of the tasks at hand. However, most LLMs lack the depth and accuracy required to understand scRNA-seq and spatial transcriptomics data analysis and the functions of related algorithms. As a result, existing tools are not yet capable of fully automating analysis.

To fill this gap, we propose CellAgent, an autonomous intelligent agent powered by GPT-4 for scRNA-seq and spatial transcriptomics data analysis. CellAgent is capable of directly understanding natural language task descriptions and autonomously completing specified tasks with high quality. By integrating “deep-thinking” reasoning with automated execution for sophisticated workflows, CellAgent systematically decomposes complex tasks into manageable steps, intelligently selects and executes appropriate tools, and iteratively refines outcomes through self-reflective optimization. Our main contributions are four-fold. Firstly, we propose a multi-agent hierarchical decision-making framework designed to simulate a “deep-thinking” workflow, ensuring that each step is consistent with the completion of the overall task. Secondly, to address the limitations of LLMs in comprehending scRNA-seq and spatial transcriptomics data analysis, we curate essential expert knowledge and introduce sc-Omni, a comprehensive toolkit that integrates a wide range of tools for scRNA-seq and spatial transcriptomics data analysis. Thirdly, we introduce a self-reflective optimization mechanism that automatically and iteratively refines analysis outcomes. To support this, we develop automated evaluation methods for multiple scRNA-seq and spatial transcriptomics data analysis tasks, effectively replacing traditional manual assessments. Furthermore, to enable efficient collaboration among multiple agents, we propose a global and local integrative memory control mechanism that systematically stores and manages different types of historical information, optimizing retrieval efficiency and enhancing task execution.

Compared to traditional tools, CellAgent supports natural language interaction and enables fully automated, unattended task execution. Extensive benchmarking demonstrates that CellAgent’s performance in task completion, quality, and efficiency, is comparable to that of human experts, and in certain aspects, it even surpasses human experts. Specifically, CellAgent achieves an average execution success rate of more than 96% across over 60 datasets. Furthermore, CellAgent demonstrates competitive performance across various downstream tasks, such as cell-type annotation, batch correction, trajectory inference, spatial domain identification, and spatial transcriptomics imputation. In summary, CellAgent achieves state-of-the-art results in a dialogue-based manner, outperforming existing methods. To promote collaboration and facilitate efficient single-cell RNA sequencing and spatial transcriptomics data analysis, we have open-sourced the sc-Omni toolkit and provided an interactive online platform for CellAgent, enabling seamless integration with the broader research community.

## 2 Results

### 2.1 Overview of CellAgent

The overall framework of CellAgent is shown in Fig. 1. Given input data along with a task described in natural language, CellAgent can autonomously and intelligently perform scRNA-seq and spatial transcriptomics data analysis (Fig. 1a). CellAgent introduces a multi-agent collaboration paradigm, strategically distributing tasks among three core LLM-driven agents: Planner, Executor, and Evaluator. The Planner understands the user requirements and decomposes the complex tasks. The Executor dynamically selects appropriate tools, generates code, and executes the code. The Evaluator assesses the result without human intervention to achieve self-reflective optimization.

**Fig. 1:**
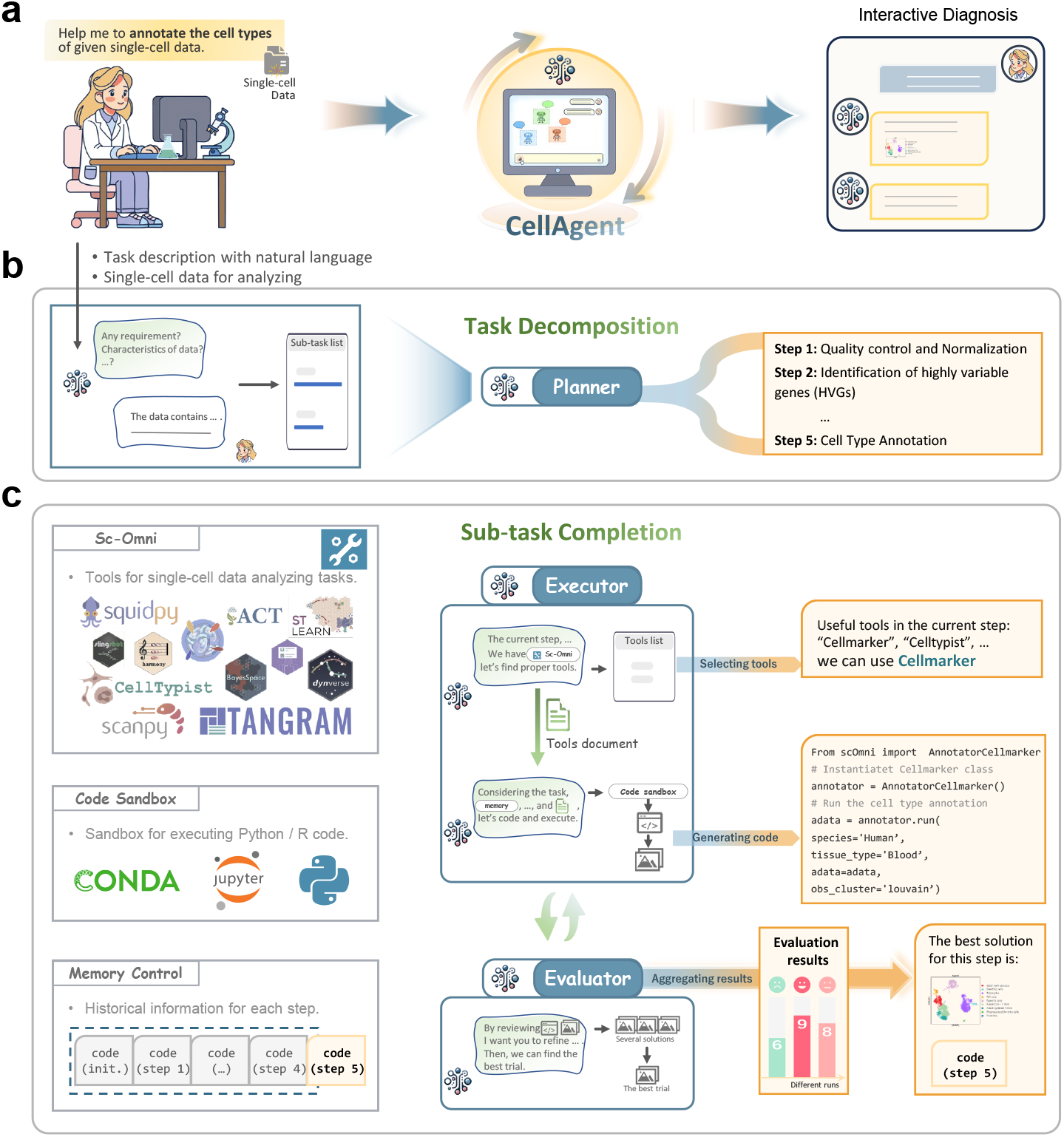
Schematic of the CellAgent Framework. **a**, Users can obtain high-quality, automated results tailored to their specific needs simply by interacting with CellAgent through natural language. **b**, CellAgent employs a hierarchical framework for task processing, consisting of high-layer planning and lower-layer execution phases. The planner is responsible for fine-grained task decomposition, based on data characteristics and relevant information. **c**, The lower-layer execution phase ensures the sequential and high-quality completion of these subtasks. During execution, the Executor selects the most suitable tools from sc-Omni, utilizing its comprehensive programming and biological knowledge to facilitate code generation and execution. Subsequently, the Evaluator rigorously assesses the execution outcomes and, if necessary, proposes optimization strategies to enhance the precision and effectiveness of the results. Through this self-reflective optimization mechanism, CellAgent ultimately synthesizes the outputs of all subtasks to generate a final result that aligns with the user’s requirements.

Inspired by the deliberate, analytical problem-solving approach of human experts, CellAgent simulates “deep-thinking” through a hierarchical multi-agent framework. Leveraging natural language comprehension, the Planner engages in high-level reasoning to systematically decompose complex tasks, performing thoughtful planning based on user-provided data and requirements (Fig. 1b and Supplementary Fig. 1). To address the limitations of LLMs in understanding biological expertise and planning subtasks, we incorporate expert-curated prompts to enhance bioinformatics sensitivity and domain-specific understanding of the Planner.

Subsequently, lower-level Executors implement the decomposed subtasks in a step-wise manner, applying in-depth knowledge of scRNA-seq and spatial transcriptomics data analysis. They iteratively refine output, referencing tool documentation to make informed adjustments and minimize execution errors (Fig. 1c and Supplementary Fig. 2). This combination of strategic planning and iterative execution embodies the essence of “deep-thinking,” enhancing both the accuracy and reliability of the analysis. To enhance the capability of the Executor in selecting appropriate tools and generating accurate code for scRNA-seq and spatial transcriptomics data analysis, we have developed a versatile and meticulously curated toolkit, sc-Omni. sc-Omni integrates tools from diverse sources, including open-source software (e.g., Scanpy, Squidpy, stLearn[6, 24, 25]), widely adopted and validated software libraries (e.g., NumPy, Pandas[26, 27]), and well-established algorithms published in top-tier journals (e.g., Tangram, scVI [28, 29]). Covering a broad spectrum of tools, sc-Omni supports various downstream tasks in scRNA-seq and spatial transcriptomics data analysis, including batch effect correction, cell type annotation, trajectory inference, spatial domain identification, and spatial transcriptomics imputation, among others (Table 1). Moreover, sc-Omni is accompanied by comprehensive documentation, featuring detailed annotations on tool usage and parameter settings, which allows Executor to learn the functions of these tools. The hallmark of sc-Omni lies in its diversity and flexibility, enabling Executor to adapt tool selection and generate available code based on specific user requirements and data characteristics.

**Table 1:** The overview of the sc-Omni toolkit for single-cell RNA sequencing and spatial transcriptomics data analysis.

| Task Name | Task Category | Tools Involved | Programming Language |
| --- | --- | --- | --- |
| Removal of Low-Quality Cells and Genes | Preliminary Analysis | scanpy | Python |
| Expression Normalization | Preliminary Analysis | scanpy | Python |
| Identification of Highly Variable Genes | Essential Analysis | scanpy | Python |
| Dimensionality Reduction | Essential Analysis | PCA_scanpy, UMAP_scanpy, t-SNE_scanpy, etc. | Python |
| Neighbor Graph Computation | Essential Analysis | scanpy, squidpy | Python |
| Clustering | Essential Analysis | louvain_scanpy, leiden_scanpy, KMeans_scanpy, etc | Python |
| Cell Type Annotation | Advanced Analysis | Celltypist, ScType, CellMarker2.0, etc. | Python |
| Differentially Expressed Gene Identification | Advanced Analysis | wilcoxon_scanpy, logreg_scanpy, t-test_scanpy, etc. | Python |
| Batch Correction | Advanced Analysis | Harmony, scVI, Scanorama, etc. | Python, R |
| Trajectory Inference | Advanced Analysis | PAGA_Dyno, SlingShot_Dyno, Scorpius_Dyno, etc. | Python, R |
| Imputation for Spatial Transcriptomics | Advanced Analysis | Tangram, SpaGE, novoSpaRc, etc. | Python, R |
| Spatial Trajectory Inference | Advanced Analysis | stLearn | Python |
| Identification of Spatially Variable Genes | Essential Analysis | squidpy | Python |
| Compute Ripley's Statistics | Essential Analysis | squidpy | Python |
| Spatial Domain Identification | Advanced Analysis | DeepST, SEDR, SpaGCN, etc. | Python, R |
| Co-occurrence Probability | Essential Analysis | squidpy | Python |
| Neighborhood Enrichment | Essential Analysis | squidpy | Python |

To ensure the reliability of outputs, CellAgent incorporates a self-reflective optimization mechanism based on feedback from the Evaluator through self-assessment and critical evaluation of intermediate outcomes (Fig.1c and Supplementary Fig. 4). This self-reflective process enables the Executor to identify potential errors, adjust hyperparameters, and select alternative tools when necessary, fostering an adaptive learning loop that improves performance over time. A key innovation of CellAgent lies in its pioneering adoption of self-reflective optimization for automated evaluation. Unlike traditional bioinformatics pipelines that rely on static workflows or predefined heuristics, CellAgent dynamically assesses its own performance, enabling iterative improvements without human intervention. Specifically, the Evaluator integrates multimodal large language models (MLLMs) for the automated assessment of three tasks, including batch effect correction, trajectory inference, and spatial domain identification. For cell-type annotation, the Evaluator leverages GPT-4 to aggregate cell-type annotation results from multiple tools. For spatial transcriptomics imputation, the Evaluator assesses the performance of various imputation methods by predicting the expression of known genes, identifying the most effective approach. Then, the Executor applies this optimal method to infer the expression of unknown genes, enabling robust downstream analysis. In summary, LLM-driven automated assessment replaces human judgment in CellAgent, enabling it to simulate human-like critical thinking and decision-making.

### 2.2 CellAgent enables natural language interactive data analysis efficiently

CellAgent proposed a natural language interactive framework to analyze the scRNA-seq and spatial transcriptomics data efficiently. To demonstrate how to use CellAgent for scRNA-seq data analysis, we provide an illustrative example of clustering task on a PBMC dataset [3] (Fig. 2). After taking the input data and task description from the user, CellAgent initially establishes a comprehensive plan by decomposing the task into five key sub-steps, including quality control, normalization, identification of highly variable genes, dimensionality reduction, and clustering. Each sub-step is then completed progressively, and the analysis results are returned in a dialogue format. For each sub-step, the user can ask CellAgent to run this sub-step again with specific requirements by natural language interaction. Furthermore, we present an illustrative example of neighborhood analysis and spatially variable gene identification on a mouse brain dataset [6]. In this example, we begin at the point where the preprocessing and clustering steps have already been completed. Leveraging the capabilities of the Planner, CellAgent effectively decouples the two input tasks, performing the analysis and visualizing the results separately (Fig. 3).

**Fig. 2:**
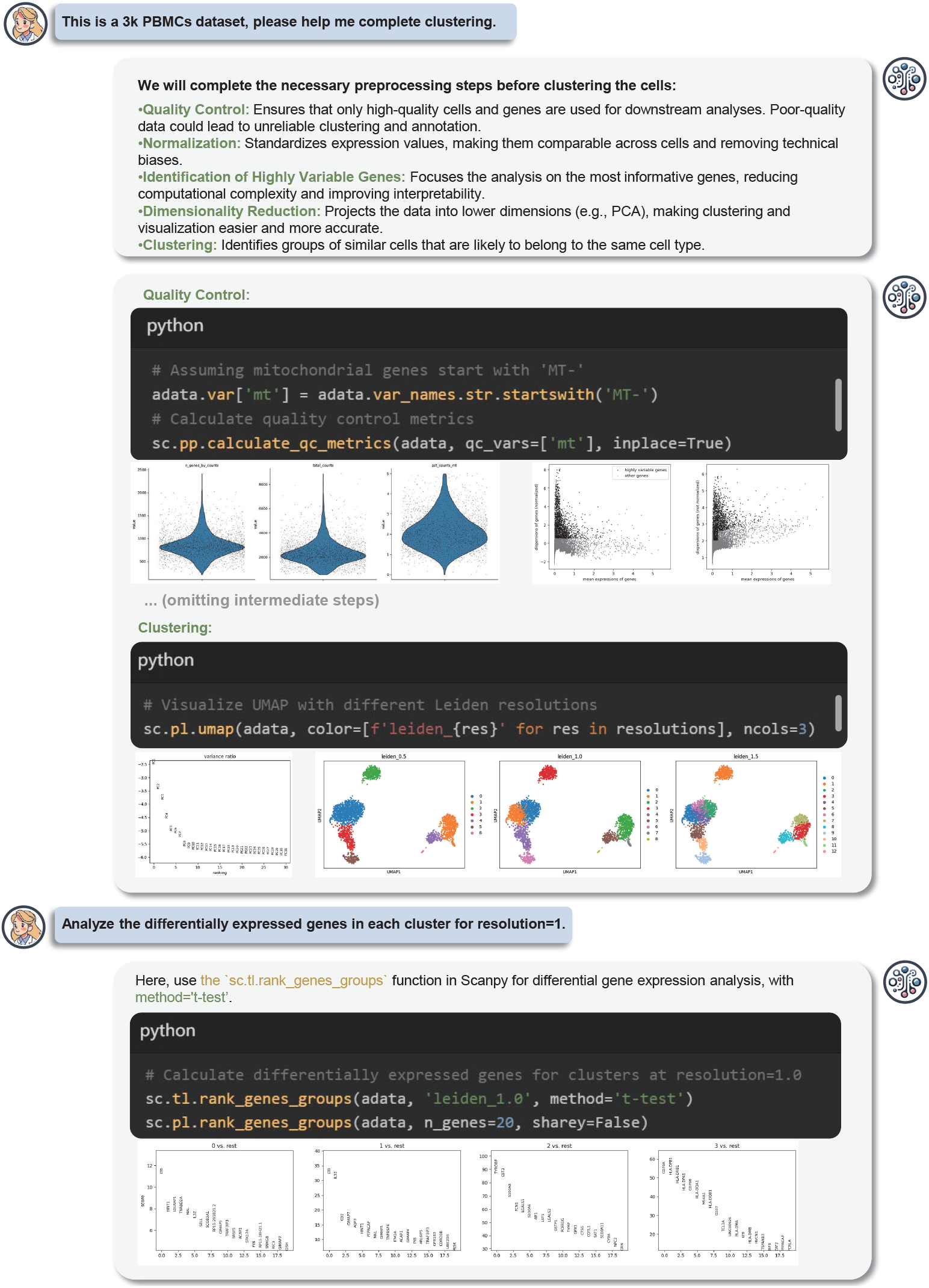
Example of using CellAgent for single-cell transcriptomics data analysis. In this case, the user provided a 3k PBMC dataset for clustering and differential gene expression analysis. CellAgent performs task planning, executes code, and generates visual results. Finally, the analysis results are delivered back to the user.

**Fig. 3:**
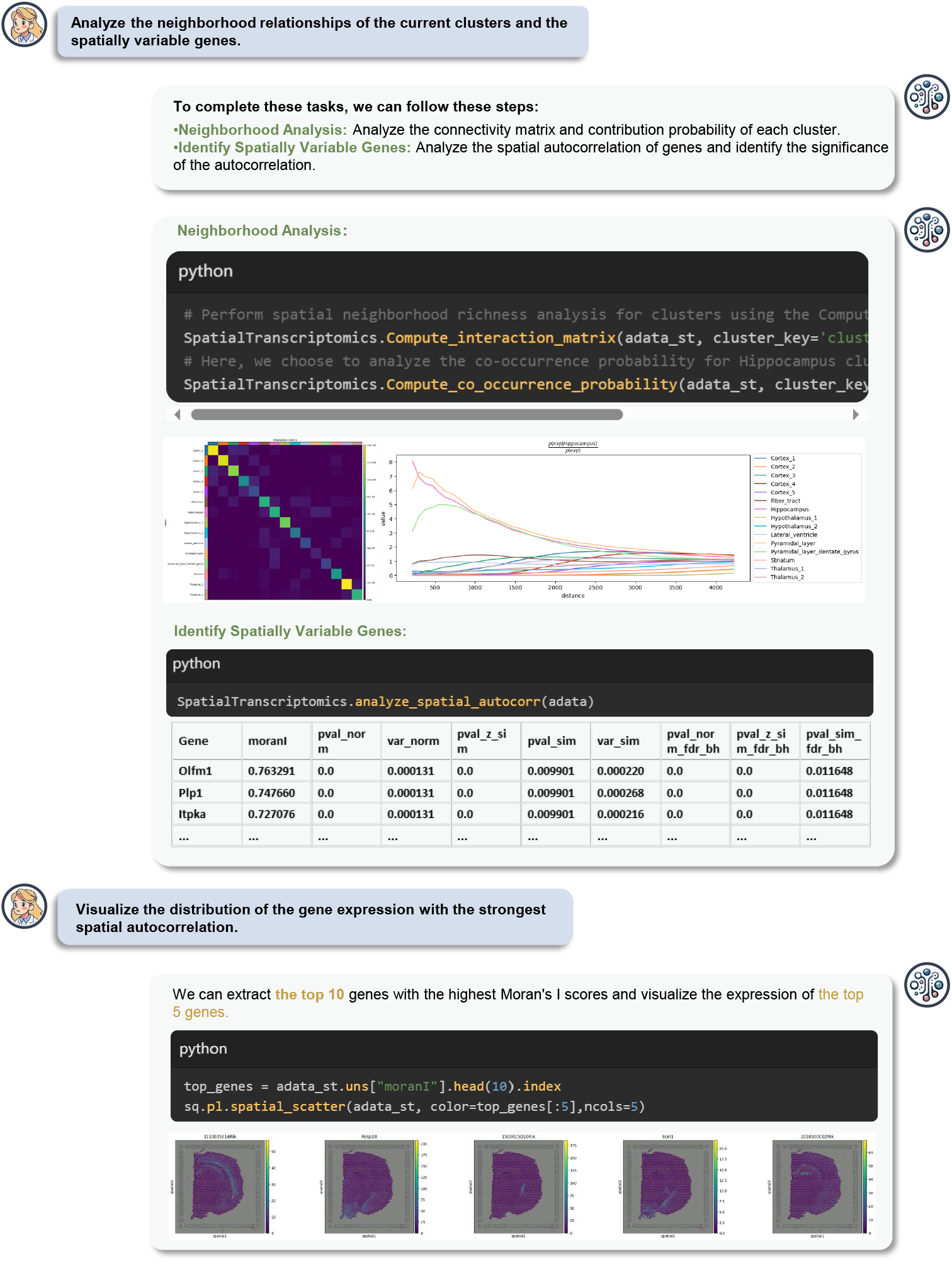
Examples of using CellAgent for spatial transcriptomics data analysis. This example encompasses comprehensive spatial transcriptomics data analysis and visualization on a 10X Visium mouse brain dataset, including spatial interaction matrix analysis, co-occurrence probability of each cluster, and identifying spatially variable genes.

To comprehensively evaluate CellAgent in data analysis tasks, we designed a benchmark consisting of eight major tasks of scRNA-seq and spatial transcriptomics analyses. These tasks include preprocessing and clustering, cell type annotation, batch effect correction, trajectory inference, spatial transcriptomics imputation, spatial domain identification, spatially variable gene identification, and spatial neighborhood analysis (Supplementary Table 1). More than 60 datasets are involved in this benchmark (Supplementary Table 8). These datasets cover different sequencing technologies, different types of tissues, and different amounts of cells.

First, we compare the ability of CellAgent with human experts. Aforementioned tasks were independently conducted by both human experts and CellAgent. Five human experts are hired for each task. Efficiency and quality are used as evaluation metrics. For efficiency, we compare the time cost of human experts and CellAgent to complete each task. For quality, five evaluators scored the results generated independently by human experts and CellAgent from 0 to 10 based on processing logic, produced code quality, and result presentation, where higher scores indicate better quality. In the efficiency evaluation, CellAgent, with an average time cost of 8 minutes, outperformed human experts with the time cost of 13 minutes (Fig. 4a). Specifically, CellAgent demonstrated advantages in complex tasks, such as spatial transcriptomics imputation, which require proficient programming skills. In the quality evaluation, CellAgent performed comparably with human experts in eight tasks, with an average quality score of 0.25 higher than the score of human experts (Fig. 4b). Then, for each task, we use GPT-4 to generate code and compare its execution success rate with that of CellAgent. The success rates of CellAgent in these tasks are significantly higher than GPT-4, which is only 23.875% on average. Noted that CellAgent@3, which supports up to three iterations of self-debugging based on error messages, achieved an average success rate of 96% (Fig. 4c).

**Fig. 4:**
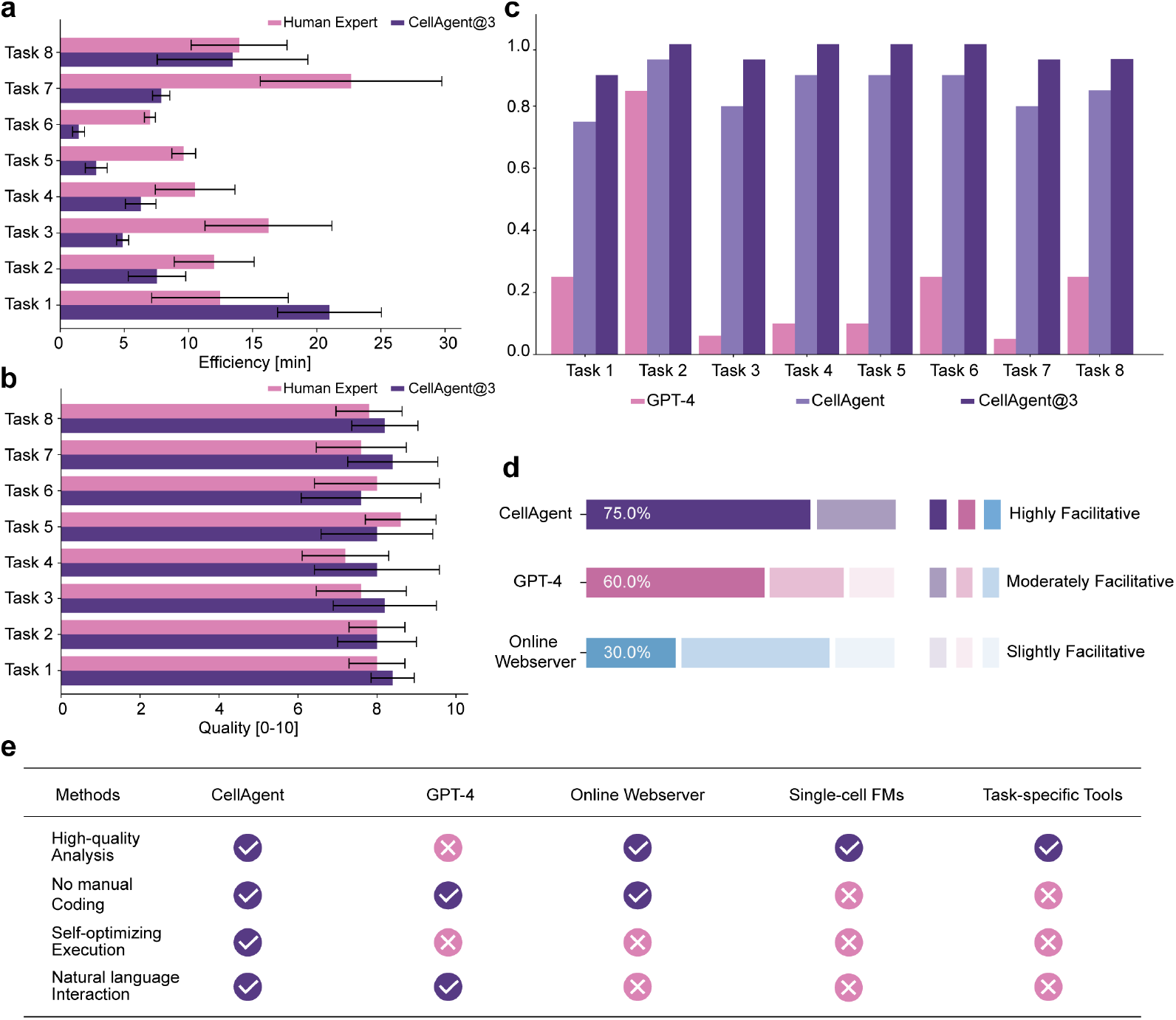
Performance comparison of CellAgent with other methods. **a**, Comparison of the efficiency of CellAgent and Human Expert on eight tasks involving scRNA-seq and spatial transcriptomics data analysis. Efficiency was assessed in minutes. **b**, Comparison of the quality of CellAgent and Human Expert on eight tasks involving scRNA-seq and spatial transcriptomics data analysis. Quality was rated on a scale from 0 to 10, with evaluations conducted by evaluators (n=5). **c**, Comparison of the success rates of GPT-4, CellAgent, and CellAgent@3 (which supports up to three iterations of self-debugging based on error messages) across the 8 tasks. **d**, Assessment of the facilitation of CellAgent, GPT-4, and Online Webserver on scRNA-seq and data analysis, as evaluated by 20 participants. **e**, Comprehensive comparison of CellAgent, GPT-4, online web servers, single-cell foundation models (single-cell FMs), and task-specific tools.

Furthermore, we assess the usability of CellAgent and other programming-free tools, including GPT-4 and an online web server [30]. The assessment involves 20 participants, comprising ten experts and ten individuals with no prior experience in biological data analysis. Each participant is tasked with performing scRNA-seq and spatial transcriptomics data analysis and subsequently rating each tool as “highly facilitative”, “moderately facilitative”, or “slightly facilitative”. The results indicate that 75% of participants suggest CellAgent highly facilitative (Fig. 4d), surpassing GPT-4 (60%) and the online web server (30%).

In summary, compared to various computational tools, including online web servers, single-cell foundation models (single-cell FMs), task-specific tools, and GPT-4, CellAgent offers high-quality analysis, no manual coding, self-optimizing execution, and natural language interaction (Fig. 4e). Together, these features demonstrate the potential of CellAgent in driving progress in scRNA-seq and spatial transcriptomics data analysis.

### 2.3 CellAgent enables efficient batch correction

Batch correction aims to adjust for technical variation across different batches of scRNA-seq data, ensuring that technical discrepancies between experiments or platforms do not distort biological signals. Proper handling of batched data is thus paramount for successful and reproducible research [31]. To evaluate the performance of CellAgent in batch correction for single-cell transcriptomics data, we compared CellAgent with scVI [7], Scanorama [32], Harmony [33], CellPLM [34], scGPT [10], and Combat [35] on five datasets [36]. These datasets encompass major tissues and organs, such as the lung, pancreas, perirhinal cortex (Supplementary Table 2). Additionally, they vary in the number of cells, number of genes, and batch origins, providing a diverse and comprehensive benchmark for batch correction evaluation. We applied ten metrics to evaluate the ability of different methods to remove batch effects while preserving biological variation. Specifically, Graph Connectivity [37], PCR Comparison [37], *iLISI*_*Graph*_ [38], kBET [39], and *ASW*_*batch*_ [40] are for the removal of batch effects (batch correction). Isolated Labels [38], ARI [41], NMI [42], *ASW*_*cell*_ [40], and *cLISI*_*Graph*_ [37] are for the conservation of biological variance (bio-conservation). The overall score is derived as a weighted sum of these ten metrics and serves as the final evaluation measure [37].

The results indicate that CellAgent outperformed other methods across all datasets, achieving the highest overall score (0.67) and individual scores. Specifically, it attained a batch correction score of 0.67 and a bio-conservation score of 0.66, demonstrating its superior performance in both batch effect removal and preservation of biological variation (Fig. 5a). We also evaluate the impact of dataset variability on algorithm performance (Supplementary Fig. 7). The results demonstrate that the overall distribution of CellAgent is positioned at a higher range compared to other methods, indicating consistently superior performance across datasets. Furthermore, we visualized the cell clustering results on the human pancreas dataset after batch correction using CellAgent (Supplementary Fig. 8 and Supplementary Fig. 9). The results demonstrate the superior ability of CellAgent to align cells of the same type across different batches while preserving a distinct separation between different cell types. This is particularly evident in the clustering of alpha cells, beta cells, and ductal cells. Overall, all these results demonstrated the strength of CellAgent in batch correction for single-cell transcriptomics data.

**Fig. 5:**
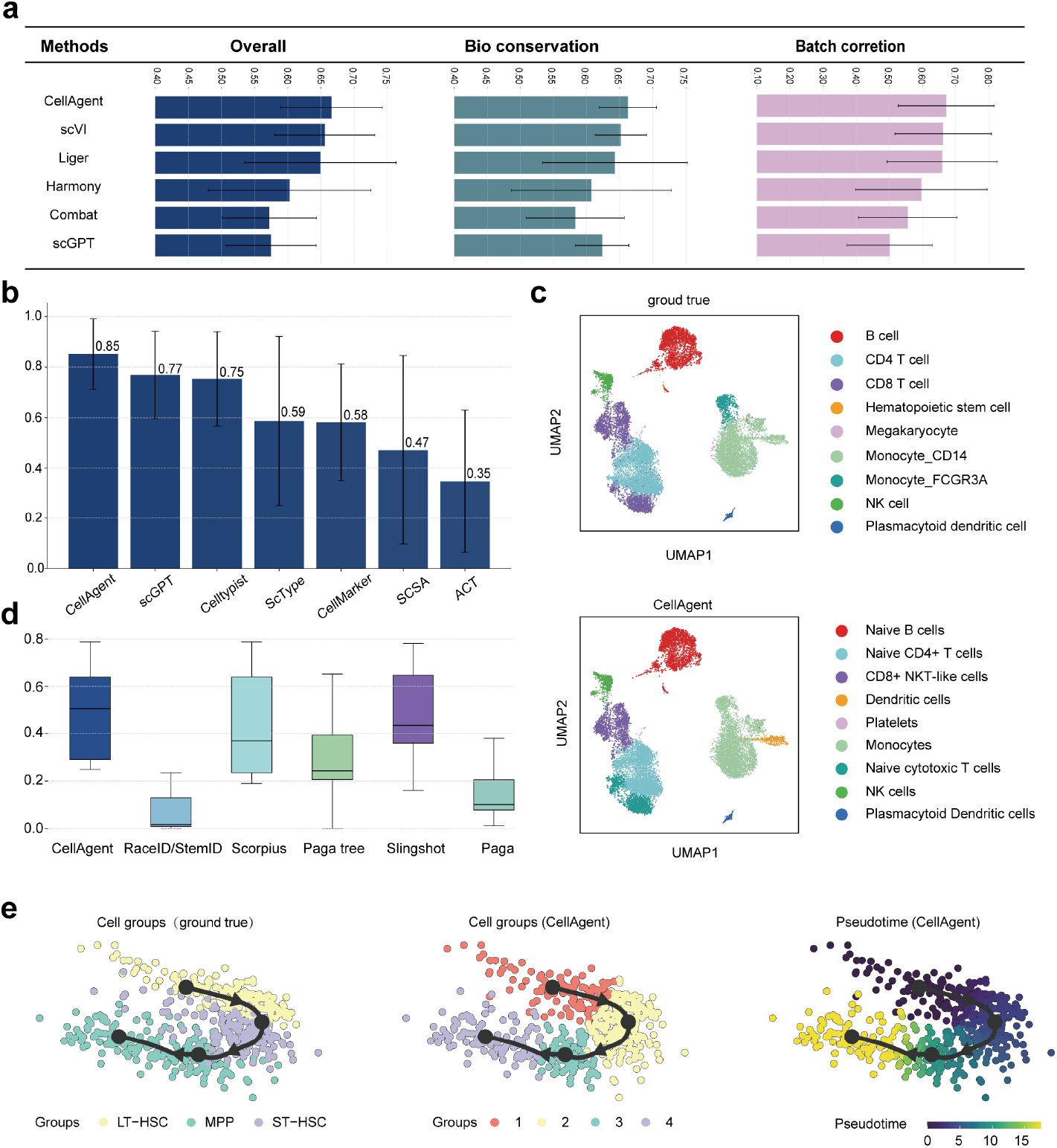
Comparison of CellAgent with other methods in single-cell transcriptomics analysis. **a**, The performance of CellAgent and other integration methods on batch correction, biology conservation, and overall score. The bar charts show the average performance across five datasets of different methods. **b**, The barplot of mean accuracy comparison between CellAgent and other cell-type annotation methods on six datasets. **c**, UMAP visualization of cell type annotation from the human PBMC dataset, colored according to the cell types annotated in the original study and the cell types predicted by CellAgent. **d**, Comparison of the performance of CellAgent and other trajectory inference methods across eight different datasets. **e**, The trajectory UMAP plot of original cell grouping, reconstructed cell grouping based on trajectory milestones by CellAgent, and pseudotime for the Aging HSC Kowalczyk dataset, going from LT-HSC, ST-HSC to MPP.

### 2.4 CellAgent facilitates more accurate cell type annotation

To evaluate the effectiveness of CellAgent in cell type annotation, we compared the results of CellAgent and other methods, such as scGPT [10], CellMarker [43], and CellTypist [44], among others, against the manual annotations provided by the original studies across six datasets. These datasets encompass various sequencing technologies, including Smart-seq2, 10X Chromium, and Drop-seq, while also spanning diverse tissues and organs, such as human peripheral blood mononuclear cells, liver, lung, and pancreas (Supplementary Table 3). Manual and automated cell type annotations were assigned specific cell ontology (CL) and general cell type names in each dataset. Annotations were categorized as “fully match” if the manual and automated annotations shared the same term or available CL cell ontology name; “partially match” if the annotations had the same or hierarchical general cell type name (e.g., fibroblasts and stromal cells) but differed in the CL names; and “mismatch” if both the general cell type names and CL cell ontology terms were different. For comparison, consistency scores of 1, 0.5, and 0 were assigned for “fully match,” “partially match” and “mismatch” respectively. The average consistency score was then calculated for each dataset across cell types.

The result shows the annotations of CellAgent fully or partially match manual annotations in an average of 85% of cell types in all datasets (Fig. 5b). This demonstrates that the cell type annotations generated by CellAgent align closely with experts’ original cell type annotations. Moreover, the visualization of cell type annotation results on the human PBMC dataset shows that the annotations of CellAgent are highly consistent with the original cell types and refined the classification of certain cell types (Fig. 5c). Specifically, CellAgent successfully differentiated CD8+ T cells in the original annotations into Naive cytotoxic T cells and CD8+ T cells, as Naive cytotoxic T cells represent a distinct subset of CD8+ T cells that have not yet encountered their specific antigen and remain undifferentiated into functional effector cytotoxic T cells [45]. Meanwhile, CellAgent also provides the differential expression and visualization of marker genes across different clusters, enabling users to gain an intuitive understanding of the characteristics of each cell population and uncover potential biological differences (Supplementary Fig. 5). These results demonstrate that by integrating multiple tools and automatically selecting optimal parameters, CellAgent provides precise annotations and enhances the understanding of cell population characteristics.

### 2.5 CellAgent enhances the performance of trajectory inference

Trajectory inference aims to determine the pattern of a dynamic process experienced by cells and model cell state transitions [46]. To assess the performance of CellAgent in trajectory inference, we compared it against five established trajectory inference methods: RaceID/StemID [47], Scorpius [48], Paga, Paga Tree [49], and Slingshot [50]. The evaluation was conducted on eight benchmark datasets with gold-standard trajectories information (supplementary Table 4) [51]. Following the previous study [51], we utilized four distinct types of metrics to evaluate different aspects of trajectories.

These metrics include: the Correlation between geodesic distances (*cor* _dist_) calculating the Spearman rank correlation between the distances, the F1 score between branch assignments (*F1* _*branches*_) measuring the harmonic mean between Recovery and Relevance of trajectories, the correlation between important features (*wcor* _features_) assessing whether the same differentially expressed features are found using the predicted trajectory as in the known trajectory, the edit distance between two trajectory topologies (*edgeflip*) quantifying the similarity in the topology between two trajectories. The overall score is calculated as a weighted average of these metrics, serving as the comprehensive evaluation of trajectory inference performance.

CellAgent demonstrated superior performance across all eight datasets, achieving the highest overall trajectory inference score of 0.496, outperforming all compared methods (Fig. 5d). Notably, in the “Aging HSC Kowalczyk” dataset, CellAgent successfully reconstructed the developmental trajectory from long-term hematopoietic stem cells (LT-HSC) to short-term hematopoietic stem cells (ST-HSC), and ultimately to multipotent progenitors (MPP) (Fig. 5e). This aligns with existing studies that describe a gradual transition from hematopoietic stem cells with high proliferative and renewal potential to those with a progressive loss of such potential, but a gain of differentiated features [52]. Beyond accurately depicting the developmental trajectories of cells, CellAgent also generated a heatmap displaying the top 20 differentially expressed genes based on their changes along the trajectory (Supplementary Fig. 6). This visualization provides an intuitive representation of gene expression dynamics across different developmental stages and identifies critical genes that may contribute to the regulation of cell fate determination. Specifically, CellAgent captured the gradual upregulation of CD48 and MPO gene expression during the hematopoietic stem cell differentiation process, consistent with their known roles in immune cell maturation and myeloid differentiation [53]. The finding supports the potential of CellAgent in identifying key genes that play critical roles in cell fate determination. These results highlight the effectiveness of CellAgent as a powerful tool for both trajectory inference and identification of key regulatory genes.

### 2.6 CellAgent enhances the performance of spatial domain identification

Spatial domain identification aims to delineate distinct regions within tissues that exhibit spatial coherence based on spatially resolved transcriptomic data [54]. To evaluate the effectiveness of CellAgent in spatial transcriptomics domain identification, we compared it with five state-of-the-art domain identification methods, including BayesSpace [55], stLearn [25], SEDR [56], SpaGCN [54], and DeepST [57] using 12 slice datasets from the dorsolateral prefrontal cortex (DLPFC) initially annotated by Maynard et al. [58]. The manual annotations based on gene expression markers and cellular morphology, delineated the cortical layers (L1–L6) and white matter (WM) regions. To quantify the accuracy of spatial domain identification, we assessed the similarity between the ground truth and the spatial domains predicted by different methods, using the adjusted rand index (ARI) as the evaluation metric [57].

CellAgent achieved the highest ARI score (0.47) across all datasets (Fig. 6a), outperforming other advanced spatial domain identification methods. Additionally, its interquartile range (IQR) was smaller, demonstrating its robustness and less susceptibility to variations in the dataset. This result demonstrates the superior capability of CellAgent in identifying spatial domains. To further validate the effectiveness of CellAgent in spatial domain identification, we compared the spatial domains predicted by different methods with the ground truth in slice 151673 (3639 spots and 33538 genes). The results revealed that CellAgent captured the intricate spatial organization of the tissue, including the differentiation of cortical layers and the white matter regions (Fig. 6b). The spatial domains identified by CellAgent were smoother and more consistent with the actual biological structure (Supplementary Fig. 14 and 15). In contrast, other methods often produced more fragmented or less biologically relevant results, which further highlights CellAgent’s ability to accurately represent complex tissue architectures and its potential for uncovering intricate spatial patterns within transcriptomic data.

**Fig. 6:**
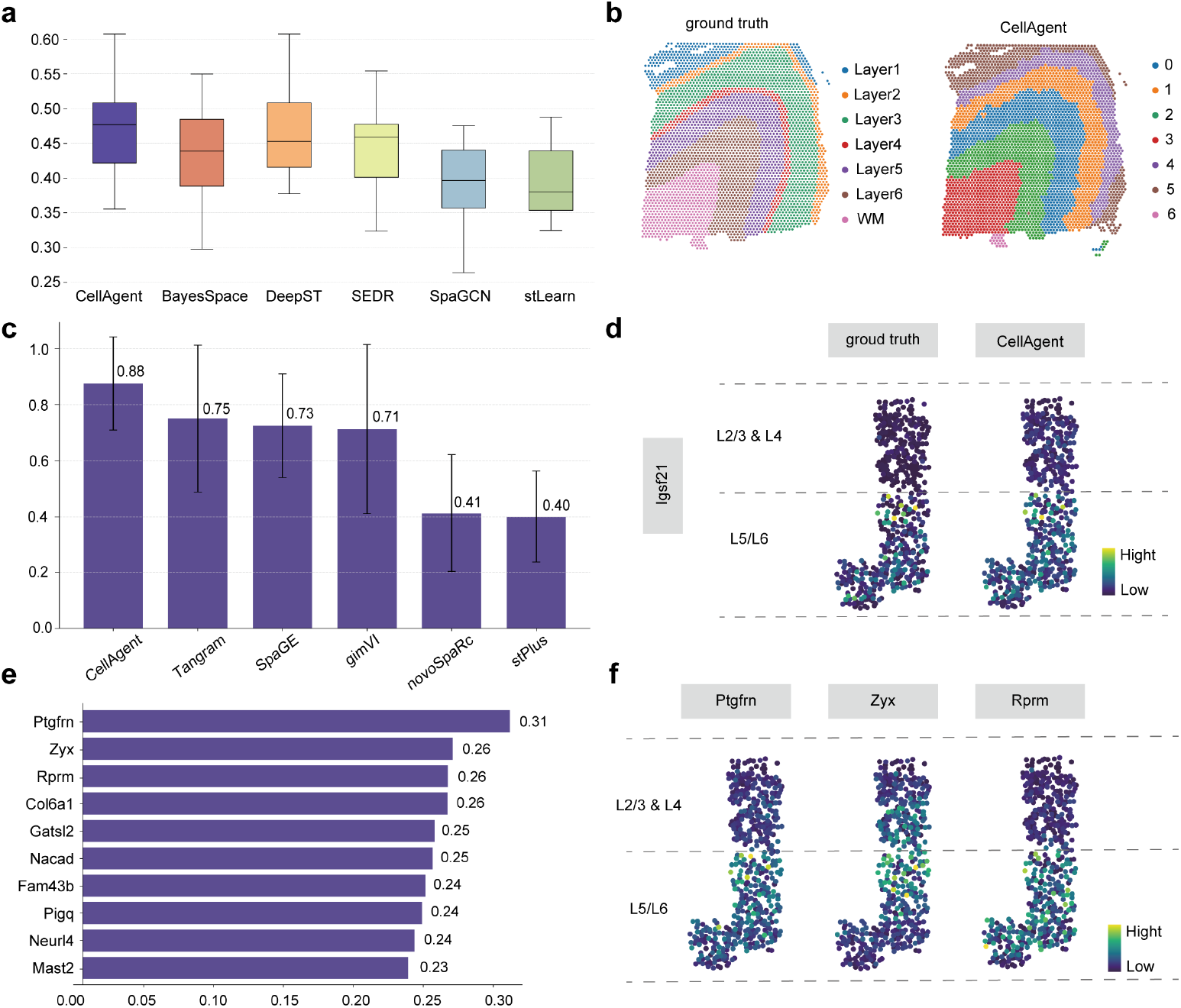
The results of CellAgent with other methods in spatial transcriptomics analysis. **a**, Boxplot of adjusted rand index (ARI) scores comparing CellAgent with other domain identification methods for the DLPFC across 12 slices. In the boxplot, the center line denotes the median, and the box limits denote the upper and lower quartiles. **b**, Visualization of the ground truth and CellAgent-identified spatial domains in slice 151673 of the DLPFC. **c**, The barplot of AS (which is aggregated from the PCC, SSIM, RMSE, and JS values; see Methods) compared between CellAgent and other spatial imputation methods across seven paired single-cell and spatial transcriptomics datasets. **d**, The ground truth spatial distribution of the gene Glg1 and the predicted distribution by CellAgent on the mouse cortex dataset. **e**, The top ten spatially variable genes were identified after imputation by CellAgent on the mouse cortex dataset. These genes were ranked based on Moran’s I spatial autocorrelation scores. **f**, The spatial distribution of top three highly variable genes: Svil, Cacna2d3, and Nt5dc2 after imputed by CellAgent.

### 2.7 CellAgent improves imputation for spatial transcriptomics

The imputation for spatial transcriptomics focuses on modeling and estimating the missing expressions over the measured spots [59]. To evaluate the performance of CellAgent in spatial transcriptomics imputation, we compared it with five state-of-the-art methods, including Tangram [29], SpaGE [60], novaSpaRc [61], stPlus [62], and gimVI [63]. These methods were applied to seven datasets derived from multiple sequencing platforms, including 10X Visium, SlideseqV2, and seqFISH (Supplementary table 6). The comparison was designed to assess the ability of each method to accurately impute spatial transcriptomics data, taking into account the distinct technical characteristics of each platform.

We employed four different metrics to assess various aspects of imputation performance. In particular, Pearson correlation coefficient (PCC) quantifies the correlation between each gene’s predicted spatial expression profile and the ground truth. The structural similarity index measure (SSIM) assesses the similarity of scaled gene expression values. Root mean square error (RMSE) calculates the discrepancy between the ground truth and predicted spatial expression. Lastly, Jensen-Shannon divergence (JS) measures the differences in spatial distribution probabilities. To provide a comprehensive evaluation, we combined these four metrics into the Accuracy Score (AS), a weighted average of PCC, SSIM, RMSE, and JS [64].

The results show that CellAgent outperformed all other methods across the four evaluation metrics, with a RANK_PCC_ score of 0.90, RANK_SSIM_ score of 0.90, RANK_JS_ score of 0.80, and RANK_RMSE_ score of 0.90 (Supplementary Fig. 16). Furthermore, CellAgent achieved the highest AS of 0.88, outperforming the suboptimal method, Tangram, by 17% (Fig. 6c). CellAgent demonstrated superior robustness across diverse datasets and exhibited significant adaptability to various sequencing platforms compared to other methods. Specifically, on the mouse cortex dataset, the predicted expression of Igsf21, a marker gene in the mouse cortex [65], closely aligned with the true expression patterns (Fig. 6d). In addition, we conducted Moran’s I spatial auto-correlation analysis on the imputed mouse cortical spatial transcriptomics data to identify the top ten spatially variable genes (Fig. 6e). The result shows Ptgfrn, Zyx, and Rprm emerged as the highest-ranked genes. Further exploration of existing literature, we found that Rprm was predicted to be highly expressed in the L5/L6 layers of the cortex (Fig. 6f), consistent with its critical role in neuronal synaptic signaling and plasticity [65]. This discovery underscores the potential of CellAgent in capturing meaningful spatial gene expression patterns.

## 3 Discussion

Single-cell RNA sequencing and spatial transcriptomics data analysis are essential for driving biological discoveries. However, conducting high-quality data analysis requires profound domain knowledge and proficient programming skills, posing a significant challenge for researchers, and most existing tools lack support for natural language interaction and have limitations in delivering high-quality automated analysis of single-cell and spatial transcriptomics data. To address these limitations, we introduce CellAgent, an autonomous, LLM-driven framework that integrates natural language interactions, a multi-agent hierarchical decision-making system, and a self-reflective optimization mechanism to automate single-cell RNA sequencing and s data analysis. This approach simulates a “deep-thinking” workflow, systematically decomposing complex tasks, intelligently selecting tools, and iteratively refining results to achieve high-quality data analysis. Compared to single-cell foundation models, online Web-server, and other tools, CellAgent offers greater flexibility, broader task coverage, and an enhanced user experience through a natural language interactive, dialogue-based approach. Through extensive experiments on a diverse set of datasets across single-cell RNA and spatial transcriptomics data analysis tasks, CellAgent has demonstrated both competitive execution efficiency and analytical quality, highlighting its potential for automated data analysis. In summary, CellAgent is a versatile and scalable tool that reduces the complexity and cost of single-cell RNA sequencing and spatial transcriptomics data analysis. The development of CellAgent will establish a new paradigm in bioinformatics, broadening the role of generative AI in scientific discovery and potentially leading to novel insights into biological systems.

## 4 Data availability

For the batch correction task, we utilized five datasets from different tissues and organs, including lung, pancreas, perirhinal cortex, and immune. These datasets are available at https://figshare.com/articles/dataset/Benchmarking_atlas-level_data_integration_in_single-cell_genomics_-_integration_task_datasets_Immune_and_pancreas_/12420968 and https://cellxgene.cziscience.com/collections/283d65eb-dd53-496d-adb7-7570c7caa443. For the cell type annotation task, we employed the following datasets: PBMC dataset (https://www.10xgenomics.com/resources/datasets/8-k-pbm-cs-from-a-healthy-donor-2-standard-2-1-0); pancreas dataset, including Smart-seq2, 10X Chromium, and Drop-seq (https://hemberg-lab.github.io/scRNA.seq.datasets/human/pancreas/); immune dataset (https://figshare.com/ndownloader/files/25717328); and liver dataset (https://www.livercellatlas.org/index.php#datasetsIndex). For the trajectory inference task, we employed the Aging HSC Kowalczyk dataset, containing developmental trajectories of human hematopoietic stem cells; the human-embryos Petropoulos dataset, containing developmental trajectories of human embryonic cells; the NKT-differentiation Engel dataset from the thymus; and the pancreatic-alpha-cell-maturation, cellbench-SC, germline-human, cell-cycle, and stimulated-dendritic-cells-PIC datasets. These datasets are deposited on Zenodo (https://doi.org/10.5281/zenodo.1443566). For the spatial domain identification task, we used twelve slice datasets from the dorsolateral prefrontal cortex within the spatialLIBD collection (http://spatial.libd.org/spatialLIBD). For the imputation experiments, we used the following datasets: Mouse embryonic stem cell dataset (seqFISH, available at https://zenodo.org/record/3735329#.YY69HZMza3J and https://figshare.com/articles/dataset/MCADGEData/5435866); Mouse cortex dataset (seqFISH+ and Smart-seq, available at https://github.com/CaiGroup/seqFISH-PLUS and https://portal.brain-map.org/atlases-and-data/rnaseq/mouse-v1-and-alm-smart-seq); Mouse prefrontal cortex dataset (STARmap, available at https://www.starmapresources.com/data and https://www.ncbi.nlm.nih.gov/geo/query/acc.cgi?acc=GSE158450); Drosophila embryo dataset (FISH and Drop-seq, available at https://github.com/rajewsky-lab/distmap and https://www.ncbi.nlm.nih.gov/geo/query/acc.cgi?acc=GSE95025); Human breast cancer dataset (10X Visium and 10X Chromium, available at https://zenodo.org/record/4739739#.YY6NpMzaWC and https://www.ncbi.nlm.nih.gov/geo/query/acc.cgi?acc=GSE176078); Mouse kidney dataset (10X Visium and 10X Chromium, available at https://www.ncbi.nlm.nih.gov/geo/query/acc.cgi?acc=GSM5224979 and https://www.ncbi.nlm.nih.gov/geo/query/acc.cgi?acc=GSE171639); and Mouse hippocampus dataset (Slide-seqV2 and Drop-seq, available at https://singlecell.broadinstitute.org/singlecell/study/SCP815/highly-sensitive-spatial-transcriptomics-at-near-cellular-resolution-with-slide-seqv2#study-download and https://www.ncbi.nlm.nih.gov/geo/query/acc.cgi?acc=GSE116470). Details of all datasets are available in Supplementary Table 7.

## 5 Code availability

We offer an online webpage accessible at http://cell.agent4science.cn/, with all code available on GitHub after manuscript acceptance.

